# Systematic characterization of existing and novel inducible transgenic systems in human pluripotent stem cells after prolonged differentiation

**DOI:** 10.1101/2025.10.17.683097

**Authors:** Michael D. Gallagher, Andrew S. Khalil, Qi Liu, Tenzin Lungjangwa, Antar Drews, Blake F. Hanan, Moritz List, Henry Hardart, David J. Mooney, Rudolf Jaenisch

**Author notes:** equal contributions (Michael D. Gallagher, Andrew S. Khalil).

## Abstract

The ability to control transgene expression both temporally and quantitatively in human-relevant cells and tissues is a cornerstone of biomedical research. Additionally, precise transgene control is crucial for optimizing human cell-based gene therapies. Human pluripotent stem cells (hPSCs) have facilitated major advances in disease modeling and the potential for regenerative medicine. Still, they are significantly limited by the lack of inducible transgenic systems that avoid silencing but maintain robust inducibility after differentiation to defined cell lineages. Here we systematically characterize the leakiness, inducibility, and tunability of multiple existing and novel transgenic systems in hPSCs and differentiated macrophages and microglia. Notably, we report the application of a small molecule-mediated splicing switch (X^on^) that allows for tunable transgene expression both before and after differentiation, without the large protein tags required for current state-of-the-art degron-based methods. We use X^on^ to achieve tight control of reporter genes, overexpression of multiple neurodegeneration-associated genes, and Cas9-mediated genome editing. We also characterize the current limitations of this system and describe approaches that can alleviate some of these limitations. By assessing multiple transgenic systems whose inducibility spans the transcriptional, post-transcriptional, and post-translational levels, we highlight and improve upon a major technical challenge that hinders basic, translational, and clinical research in physiological human-based systems.

## INTRODUCTION

Conditional gene expression systems such as Tet-On/Off, Cre-Lox, and protein modification-based strategies provide temporally resolved, gene-specific perturbations in defined experimental contexts[1, 2]. These systems enable the timed induction or silencing of genes and their associated pathways, thereby facilitating investigations into gene function, lineage tracing, circuit analysis, and gene and cell therapies[3]. In addition to single gene studies, functional genomic approaches based on CRISPR and transposase-mediated integration allow for large-scale and genome-wide functional studies[4]. However, translating these tools to human-based platforms, particularly those involving pluripotent stem cells (hPSCs) or fully differentiated cells and tissues, remains hampered by insufficient transgene control and challenges in efficient and homogeneous transgene delivery[5, 6].

hPSCs are a uniquely enabling platform for basic and translational research due to their self-renewal and directed differentiation capacity, which enable the generation of standardized, genetically defined, and reproducible sources of virtually any human cell or tissue. Moreover, the ability to engineer hPSCs at the pluripotent stage and subsequently differentiate them into various cell types and tissues offers a powerful means to generate isogenic, experimentally tractable cells[6]. In this way, hPSC-based approaches circumvent canonical challenges with primary and terminally differentiated cells, such as heterogeneous transgene delivery and the need for repeated transfection, virus production and transduction. These advantages, combined with genetic background heterogeneity that may correlate with patient clinical phenotypes and therapeutic response[7, 8], may increase the likelihood of translating hPSC-based results into clinical use.

An often overlooked but critical limitation of current hPSC-based methods is the lack of robust inducible transgenic systems. While a few studies have reported this issue[9–11], discussions among stem cell researchers suggest a lack of awareness and frustration due to underreporting of failed experiments and wasted time and effort using systems known by others to fail. Inappropriate transgene expression before hPSC differentiation can cause unwanted cell differentiation or confound interpretation of results after differentiation is complete; conversely, transgenes must remain robustly inducible after differentiation in order to manipulate the gene(s) of interest in the appropriate cellular context[6]. Transgene dosage control is also important, as dose-dependent effects of proteins, RNAs and other molecules have been widely reported[3, 12–17]. Furthermore, continuous expression of genome perturbation effectors such as Cas9 can result in toxicity or phenotypic instability, preventing the establishment of stable hPSC lines with guide libraries[6]. The alternative option, introduction of transgenes into differentiated cell types by transfection, electroporation or viral transduction, is often toxic, inefficient, or attainable only with unwanted cellular responses[6]. As a result, study designs spanning individual gene perturbations to high-throughput screening in hPSC-based systems remain constrained relative to immortalized lines, and overcoming these hurdles is critical for advancing the experimental potential of hPSC-based systems.

Here, we have systematically characterized the strengths and limitations of several prevalent hPSC inducible expression systems. The systems examined span transcriptionally regulated (Tet-On), post-translationally regulated (degron), and post-transcriptionally regulated (X^on^) systems. We additionally evaluated new optimization strategies to improve their function, specifically in hPSC-derived myeloid cells. We chose myeloid cells (macrophages and microglia) due to their sensitive and context-dependent functions in tissue homeostasis, immune surveillance, and inflammation[18, 19], as well as a general field interest in applying functional genomic studies to better understand their roles in health and disease. Additionally, their sensitive phenotypic responses necessitate tight transgene control, and their derivation typically requires long differentiation protocols[18, 20], providing a stringent test case for this study. We highlight the X^on^ system as a novel hPSC-based system that overcomes important limitations of existing systems, and allows for tight control of reporter genes, overexpression of neurodegeneration-associated genes in microglia, and Cas9-based genome editing. Collectively, these efforts establish a foundation for developing next-generation hPSC-based gene expression platforms to advance basic and translational research.

## RESULTS

### Transposon-integrated Tet-On transgenes are not functional in hPSCs or hPSC-derived myeloid cells

We selected six candidate master microglial transcription factors (TFs) that are downregulated or silenced in cultured microglia, as reactivating these TFs may generate important insights into microglial function and improve current *in vitro* microglial cell models[21–24]. These TFs (SALL1, SMAD3, TAL1, MEF2A, MEF2C and KLF2) were cloned into validated Sleeping Beauty (SB) transposons that have been optimized for balancing robust constitutive reporter expression and tight control of the Tet-On-driven transgene[25] **(Fig. 1A)**. Three TFs were cloned into two transposon plasmids (pSBtet-GP and pSBtet-RP) separated by 2A cis-acting hydrolase element (CHYSEL) sequences that encode for ribosome skipping peptides[26, 27]. Co-transfection of each transposon with the SB100X transposase[28] into HEK293Ts **(Fig. 1B)** resulted in persistent GFP+ and tdTomato+ (tdT) cells, suggesting transposon integration. Fluorescence-activated cell sorting (FACS) isolated pure GFP+ and tdT+ populations, and doxycycline (dox) treatment robustly increased expression of the first two TFs in each transposon **(Fig. 1C–D)**, but not the third. Two bands near the expected size of TAL1 were detected in cells lacking the transgene, and did not increase in intensity in dox-treated transgenic cells **(Fig. S1A)**. KLF2 blotting revealed dox-specific high molecular weight species, potentially representing translational read-through across 2A peptides[27] **(Fig. S1B)**. HEK293Ts with stable integrations of both pSBtet-GP-TF and pSBtet-RP-TF displayed dox-inducible TF expression from both transgenes **(Fig. 1E)**.

**Figure 1:**
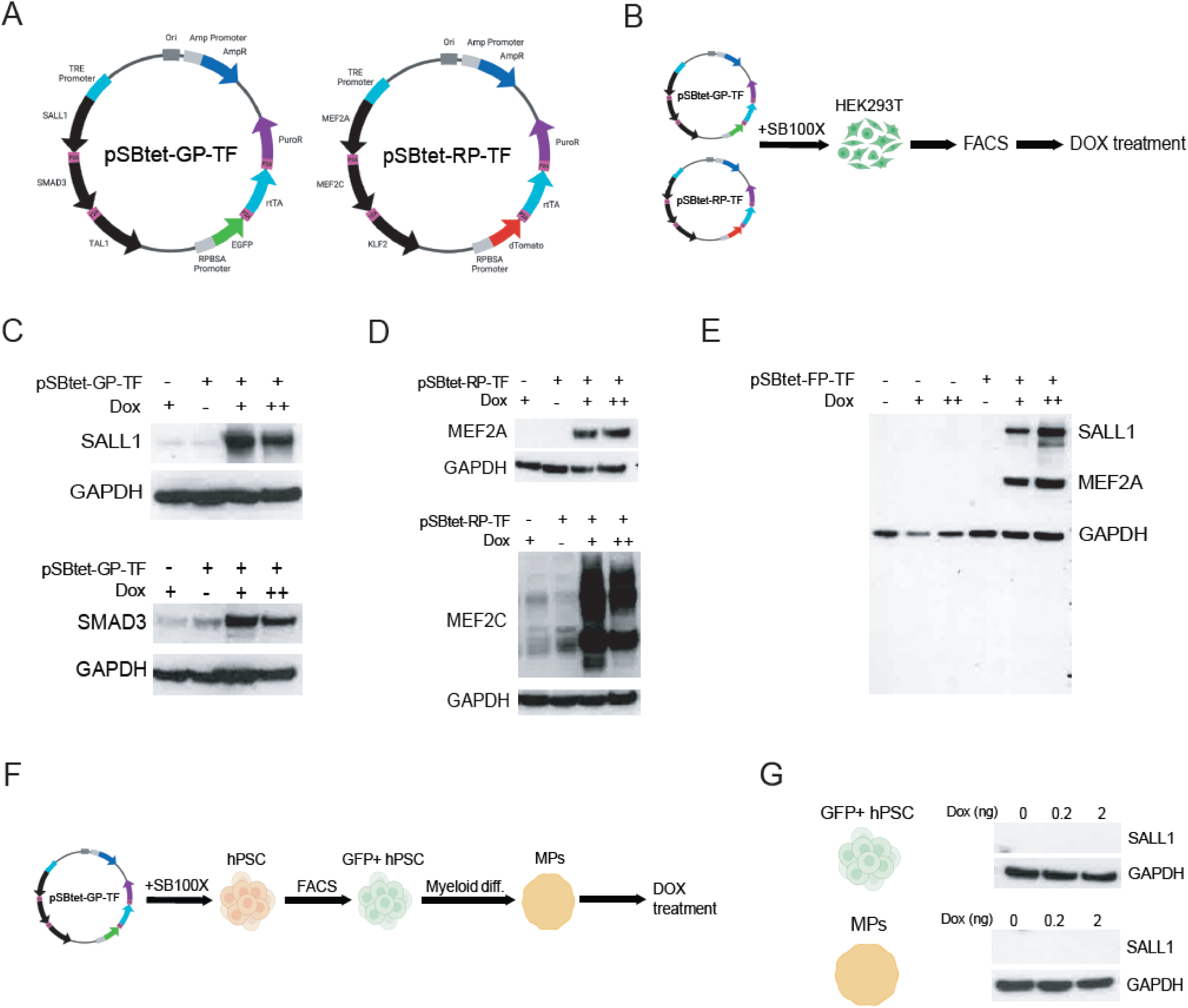
Transposon-integrated Tet-On transgenes are not functional in hPSCs or hPSC-derived myeloid cells. **(A)** Diagrams of pSBtet-GP-TF and pSBtet-RP-TF transposons for dox-inducible TF expression. **(B)** Experimental schematic of transposon testing in HEK23T cells. **(C,D)** Western blot analyses of HEK293Ts with stable integrations of pSBtet-GP-TF (panel C) and pSBtet-RP (panel D). “-“ and “+” signs indicate absence and presence of transposon and dox treatment (0.75 or 7.5ug/mL). Blotted proteins are indicated to the left of the blots. **(E)** Western blot analysis of cells with dual integrations of pSBtet-GP-TF and pSBtet-RP-TF (indicated as pSBtet-FP-TF). Expected molecular weight bands for blotted proteins are indicated to the right. **(F)** Experimental schematic of transposon testing in hPSC-derived myeloid progenitors. **(G)** Western blot analysis of SALL1 in hPSCs and hPSC-derived myeloid cells with stable pSBtet-GP-TF integrations. Blotted proteins are indicated to the right of each blot.

We then integrated the same transposons into hPSCs that we have used to generate hPSC-derived microglia-like cells (hMGLs)[22, 29] **(Fig. 1F)**. Persistent GFP+ and tdT+ cells were observed and sorted 7 days post-transfection, though at lower frequencies and with weaker fluorescence than what was observed in HEK293Ts. GFP and tdT expression diminished with passaging, and dox treatment failed to induce expression of SALL1, the first position TF, in pSBtet-GP-TF hPSCs and myeloid progenitors **(Fig. 1G)**. These myeloid cells also died within 24h of puromycin (puro) treatment (data not shown), indicating silencing of both inducible and constitutive transgenes in SB transposons.

To test whether stronger constitutive promoters might prevent transgene silencing[6, 30], we transfected hPSCs with EGFP plasmids driven by CMV, CAG and PGK promoters, and compared them to the pSBtet-GP-TF plasmid, in which EGFP is driven by a chimeric ribosomal (RPBSA) promoter[25]. CMV and PGK plasmids produced more robust GFP expression than RPBSA and CAG **(Fig. S1C)**, although CMV has been shown to be prone to silencing in hPSCs[6]. We therefore replaced the RPBSA promoter in the original pSBtet constructs with PGK, and additionally included EF1A, which is one of the strongest known promoters across diverse cell types[31] **(Fig. S1D)**. EF1A-driven plasmids (pSBtet-GP-EF1A and pSBtet-RP-EF1A) produced the brightest GFP and tdT expression, respectively, so we used them to generate stable GFP- and tdT-expressing hPSCs **(Fig. S1E–F)**. However, puro treatment killed nearly all cells in both populations, and the small number of surviving cells died 6-10 days after initiating myeloid differentiation **(Fig. S1G–H)**.

### Safe harbor-targeting does not rescue Tet-On function in hPSC-derived myeloid cells

To test whether semi-random integration of SB transposons contributed to the silencing of Tet-On-driven transgenes, we generated hPSCs with a validated Tet-On transgene[32] targeted to the AAVS1 safe harbor locus, based on approaches developed in our lab. This transgene contains a splice acceptor that allows the puromycin resistance gene to be translated from the endogenous PPP1R12C mRNA, and a CAG promoter driving the reverse tetracycline transcriptional activator (rtTA) [33, 34]. We replaced the existing transgene with EGFP (pUCM-EGFP) for simple readout of inducibility, and observed robust EGFP 48h after dox treatment **(Fig. 2A)**, which was reproducible after multiple passages and freeze/thaws. However, upon differentiation to hPSC-derived macrophages (hMacs), we observed limited-to-no ability to induce expression of EGFP **(Fig. 2A)**. Similar results were observed with hMGLs and with other AAVS1-targeted Tet-On transgenes (data not shown). Previous reports of Tet-On transgene silencing in pluripotent stem cells suggest that DNA methylation of the tetracycline-responsive element (TRE) promoter is responsible, and recruitment of TRE-binding domains fused to the murine Tet1 demethylase domain rescued transgene silencing in mouse embryonic stem cells[35, 36]. To test this approach in hPSCs, we synthesized new Tet-On donor constructs that employed rtTA and the TRE-binding domain of the tetracycline transcriptional silencer (tTS) in a single polycistronic CAG-driven cassette, with or without fusion of the TET1 demethylase domain **(Fig. 2B)**. In these systems, the tTS or tTS-TET1 fusion proteins should bind the TRE promoter in the absence of dox, and prevent DNA methylation via steric hindrance and/or demethylase activity. Upon binding dox, the tTS proteins should be evicted from the TRE promoter and replaced by rtTA, resulting in transgene expression. As a control, we synthesized a standard 3^rd^ generation Tet-On (TetO3G) transgene with rtTA alone in the same format **(Fig. 2B)**.

**Figure 2:**
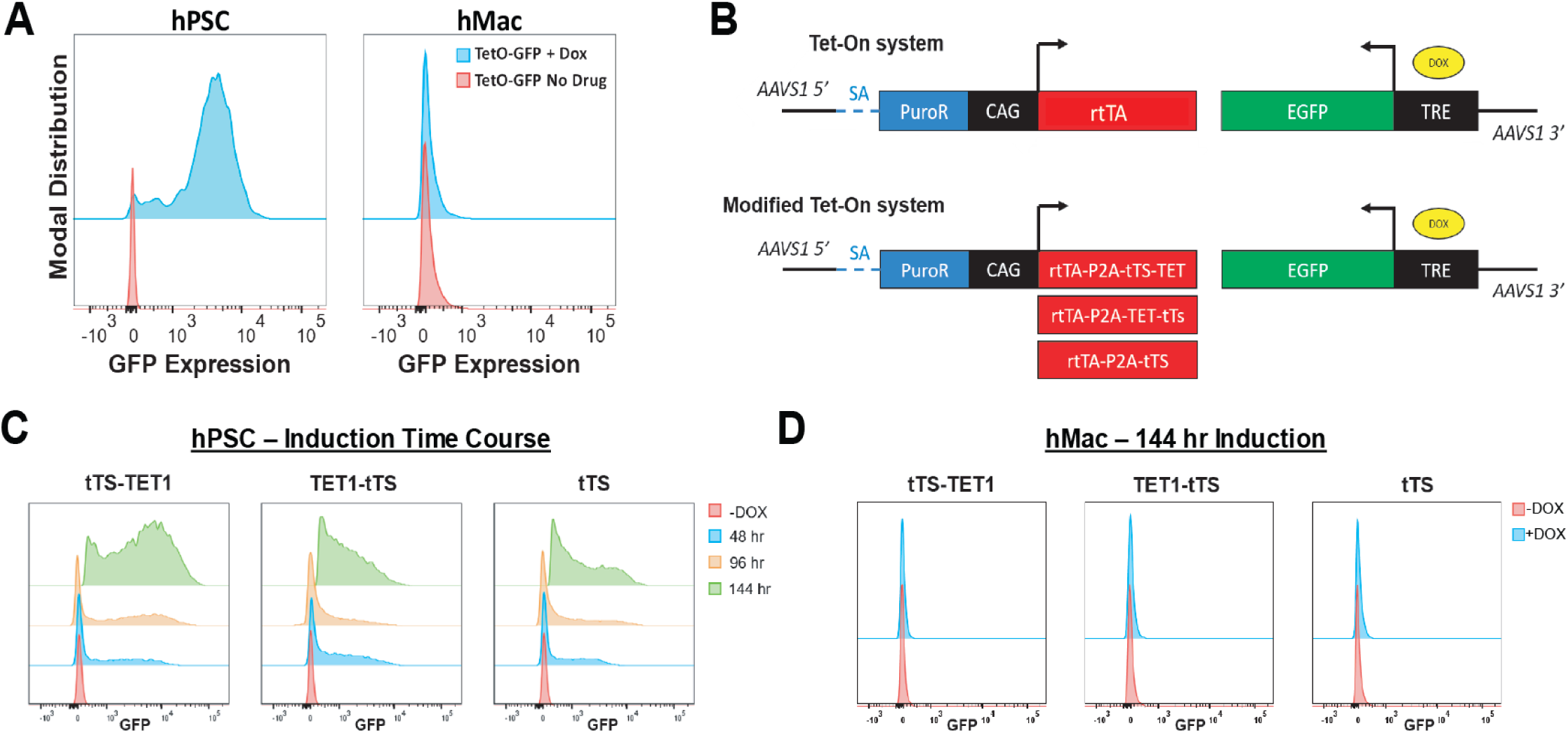
Safe harbor integration does not rescue Tet-On function in hPSC-derived myeloid cells. **(A)** Comparison of the commonly used pUCM TetO3G induction platform to drive EGFP expression in hPSCs and hMacs illustrating lack of transgene inducibility at the myeloid stage. **(B)** Schematic of the control TetO3G (top) and dual rtTA/tTS-based designs (bottom). **(C)** Representative flow cytometry plots of time-course dox-induction in rtTA-tTS designs, showing retention of inducibility in hPSCs. **(D)** Representative flow cytometry plots of time-course dox-induction in all rtTA-tTS designs, showing lack of inducibility in hMacs.

To test these constructs we performed a time course of induction in hPSCs. We observed variable but robust EGFP expression across all dual rtTA-tTS constructs, both with and without TET1 domains **(Fig. 2C)**. However, induction kinetics were delayed relative to pUCM-EGFP and our TetO3G control **(Fig. 2A, S2B)**. Among the three dual systems, the C-terminal fusion demonstrated the highest level of transgene induction, whereas the tTS alone produced the weakest induction **(Fig. 2C)**. With respect to leakiness, only the C-terminal fusion showed a marked increase in population-level green median fluorescence intensity (MFI) and percentage of EGFP+ cells, compared to non-transgenic (WT) cells **(Fig. S2A)**. However, after differentiation to hMacs, no system allowed for dox-inducible EGFP expression **(Fig. 2D)**. In summary, Tet-On systems fail to induce transgene expression after hPSC myeloid differentiation, regardless of transgene delivery method, constitutive promoter choice, targeting vector, or addition of tTS-TET1 fusion proteins.

### DHFR degron-based systems are active in hPSC-derived myeloid cells but not tightly regulated

Having observed silencing with multiple Tet-On systems and the inability to prevent silencing with TET1 demethylase fusion proteins, we tested whether other existing inducible methods allow for robust, tight transgene control in hPSC-derived myeloid cells. We selected the dihydrofolate reductase (DHFR) degrons, which target C-terminally-fused proteins for degradation by the ubiquitin-proteasome pathway (UPP), and have been used for a variety of hPSC, immortalized cell line, and in vivo applications[37]. The small molecule trimethoprim (TMP) inhibits DHFR-mediated degradation, and can be added when stable protein levels are desired **(Fig. 3B)**. We fused optimized N-terminal and C-terminal DHFR degrons[38] to a CAG-driven EGFP transgene, generating single and dual degron-tagged transgenes (**Fig. 3A)**. We additionally generated a separate dual-degron-tagged EGFP using the exact degron sequences previously used for diverse CRISPR screening applications in hPSC-derived neurons and microglia[39–42]. All constructs were targeted to the AAVS1 locus in hPSCs, as described above.

**Figure 3:**
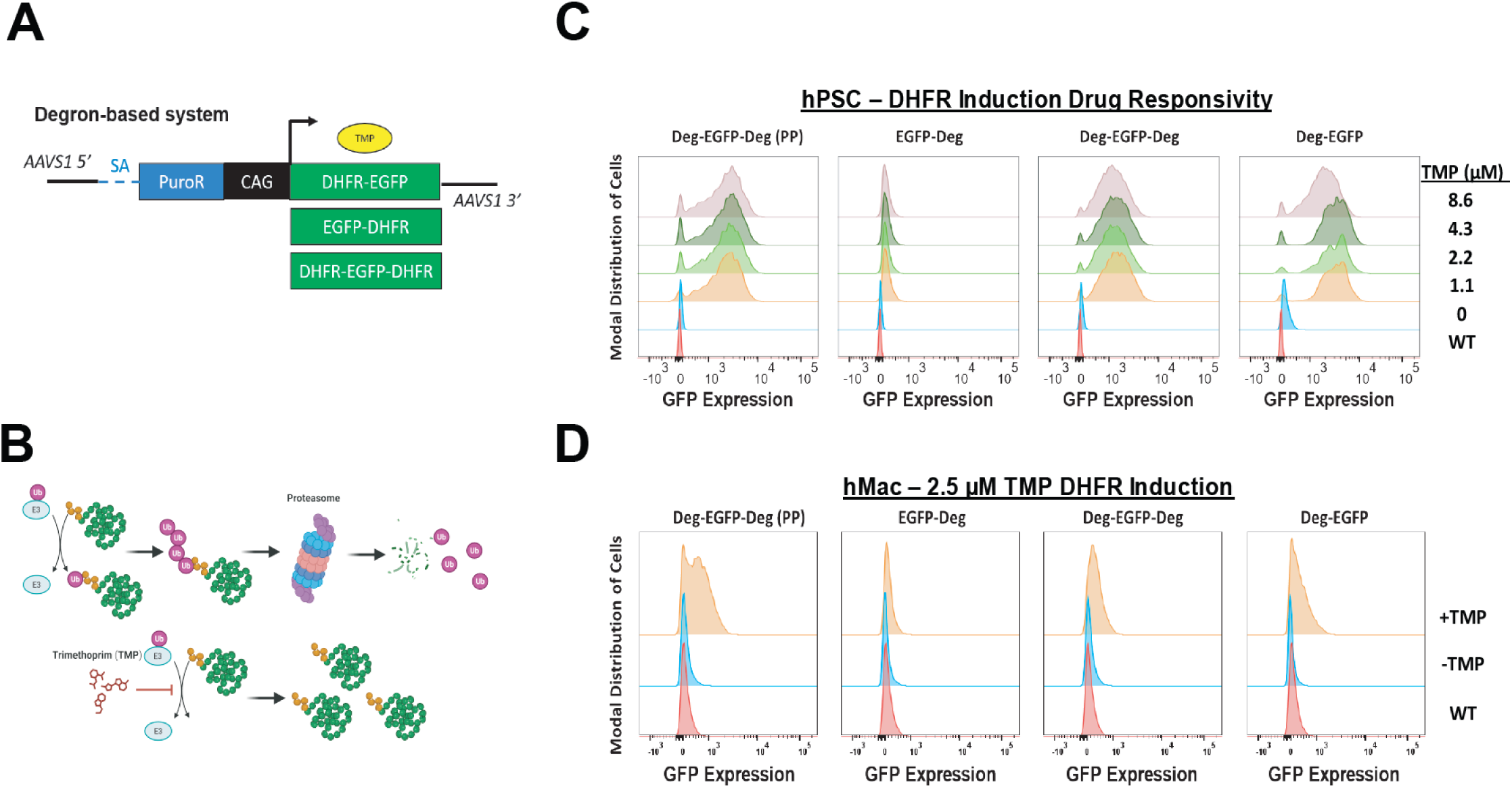
DHFR degron-based systems are functional in hPSC-derived myeloid cells but not tightly regulated. **(A)** Schematic for DHFR-tagging of N-, C-, and dual termini of EGFP in a CAG-driven format in the AAVS1 locus. **(B)** TMP-mediated stabilization of the DHFR-tagged fusion proteins to prevent degradation via the ubiquitin-proteasome pathway. **(C)** Representative flow cytometry plots of TMP dose-response induction in all DHFR degron designs, showing inducibility in hPSCs in all but the C-terminally tagged EGFP. WT represents unengineered hPSCs. **(D)** Representative flow cytometry plots of 2.5uM TMP induction in all DHFR degron designs, showing retention of inducibility in hMacs with all but the C-terminally tagged EGFP. WT represented unengineered hMacs.

To test the inducibility of the different degron designs, we performed a dose-response study of TMP and assessed EGFP by flow cytometry 24h later. We observed robust EGFP induction for the dual, N-terminal, and previously published (PP) dual degron-tagged transgenes, but not for the single C-terminal tag **(Fig. 3C)**. We did not observe tunable EGFP levels within the dose range tested, and all degrons except for the C-terminal tag were variably leaky **(Fig. S3A-B)**. To test degron performance after differentiation, we treated hMacs from each population with 2.5µM TMP, which produced the highest EGFP levels in hPSCs **(Fig. S3A)**. We observed reduced inducibility across all degrons, but similar relative performance as observed in hPSCs. Notably, substantial proportions of EGFP-cells were present in all systems **(Fig. 3D)**. In summary, these results demonstrate that DHFR degrons have limited utility in hPSC-derived myeloid cells, and do not allow for robust or tunable transgene expression.

### X^on^ inducible splicing system allows for reproducible, robust and highly tunable transgene expression

We next turned to the recently reported inducible splicing (X^on^) system[43]. X^on^ consists of a ∼1.2kb sequence from the 5’ end of the human *SF3B3* gene, and includes a pseudoexon (PE) that is only spliced into the transgene mRNA in the presence of LMI070 (branaplam), a small molecule that increases U1 snRNP recruitment to 5’ splice sites[44]. Previous studies have employed this system by moving the start codon of the gene of interest (GOI) into the PE, which results in LMI070-specific translation of the GOI with a 24 amino acid N-terminal tag[43] **(Fig. 4A)**.

**Figure 4:**
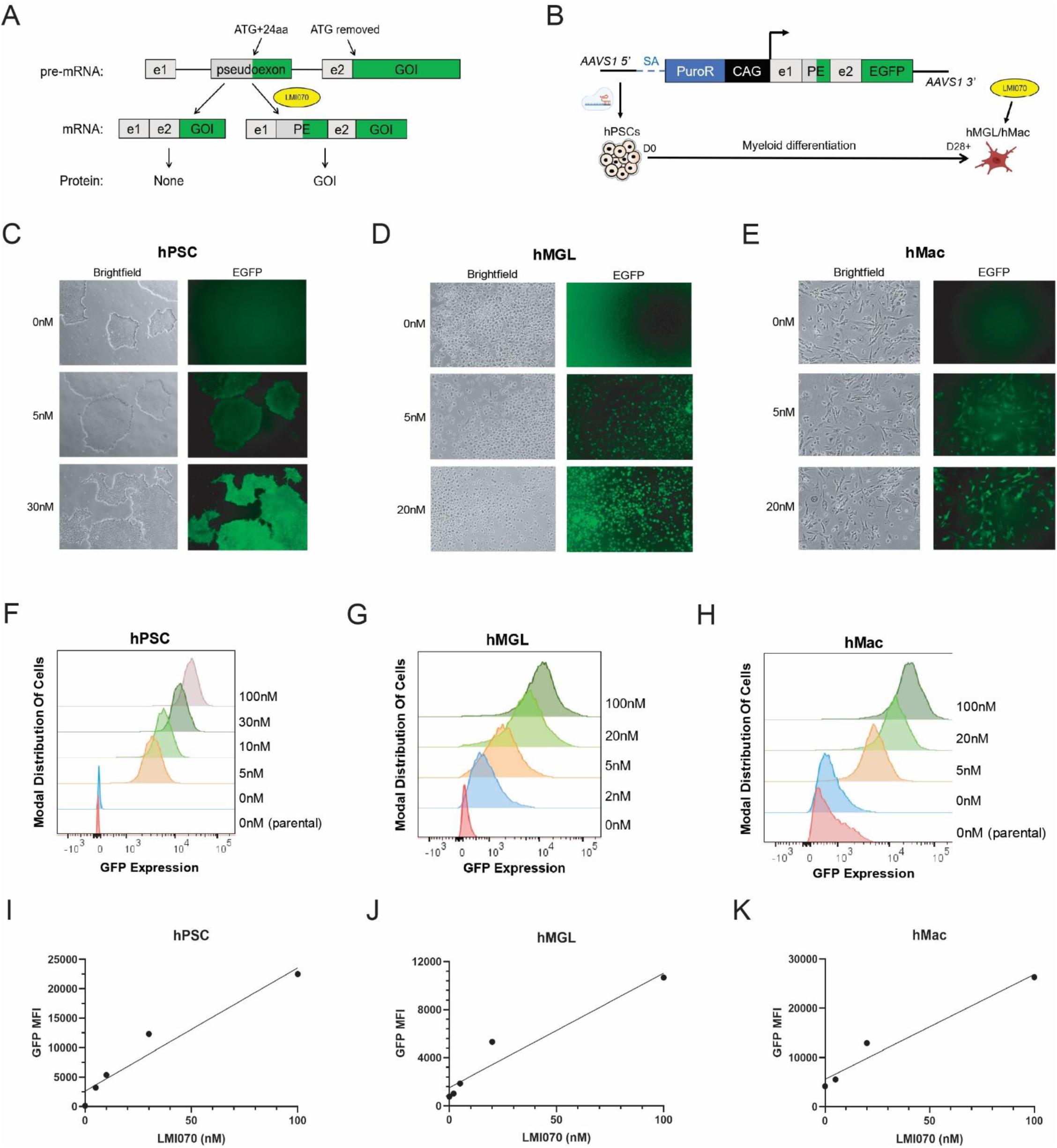
X^on^ inducible splicing system allows for reproducible, robust and highly tunable transgene expression. **(A)** Schematic of X^on^ inducible splicing mechanism. Start codon is removed from the gene of interest (GOI) so that translation requires LMI070-mediated inclusion of ATG in the pseudoexon, resulting in a 24 amino acid N-terminal tag. e1=exon 1, e2=exon 2, PE=pseudoexon. **(B)** AAVS1 targeting strategy for testing X^on^-EGFP in hMacs and hMGLs. SA=splice acceptor. PuroR=puro resistance. **(C-E)** Fluoresence microscopy images of X^on^-EGFP hPSCs, hMGLs and hMacs, treated with various doses of LMI070. **(F-H)** Quantification of EGFP by flow cytometry in dose-response experiments. “Parental” samples are WT cells that lack the X^on^-EGFP transgene. LMI070 dose is indicated to the right of each image. **(I-K)** Scatter plot representation of data from panels F-H.

To test this system in hPSCs, we targeted X^on^-regulated EGFP (X^on^-EGFP) constructs to the AAVS1 locus and differentiated them to hMacs and hMGLs using both embryoid body- and monolayer-based protocols **(Fig. 4B)**. In all cell types, X^on^-EGFP was robustly inducible and highly tunable, with expression levels strongly and linearly correlated with LMI070 dose (hPSC R^2^=0.938, P=0.0067; hMGL R^2^=0.932, R=0.0077; hMac R^2^=0.958, P=0.021) **(Fig. 4C–K, S4)**. Uniform EGFP+ hPSC populations were obtained with as low as 5nM drug dose after 24h; however, doses of 30nM or higher resulted in morphological changes reminiscent of differentiation **(Fig. 4C, F)**. For hMGLs, 5nM and 20nM doses achieved 92% and 98% GFP+ cells, respectively **(Fig. 4D, G)**. Significant autofluorescence in hMacs, consistent with a highly phagocytic monocyte-derived macrophage identity[45], made it difficult to distinguish GFP+ cells from background signal. However, >90% of cells were unambiguously GFP+ at the 20nM dose **(Fig. 4E, H)**. These results were consistent across two different hPSC backgrounds (hPSC #2 R^2^=0.941, P<0.0001) and both hMGL differentiation protocols **(Fig. S4)**. In addition, neither hMGLs or hMacs exhibited observable changes in cell morphology or apparent cell death with doses up to 100nM **(Fig. 4D-E, S4C)**. While minor EGFP leakiness was observed by flow cytometry **(Fig. 4F, S5)**, the levels observed were orders of magnitude below those seen in cells treated with even the lowest drug doses. In summary, X^on^ overcomes multiple limitations of existing inducible hPSC-based systems, and allows for highly tunable and silencing-resistant transgene expression.

### Incorporating N-terminal 2A peptides can improve inducible X^on^ systems

To test the utility of X^on^ for genes relevant to myeloid cell function, we replaced EGFP with SALL1. Western blotting revealed that X^on^-SALL1 hPSCs expressed SALL1 protein at higher levels than WT hPSCs, even in the absence of LMI070, and with no further induction after LMI070 treatment. In contrast, EGFP exhibited dose-dependent and LMI070-specific expression **(Fig. 5A)**, consistent with the flow cytometry results in **Fig. 4**. We observed similar results with hPSCs from a different background (**Fig. S6A)** and in HEK293Ts transfected with the X^on^-SALL1 donor construct **(Fig. 5B)**. To assess LMI070-mediated pre-mRNA splicing, we designed qPCR primers to detect the expected mRNA species, and compared the results with those from X^on^-EGFP hPSCs **(Fig. 5C)**. Two primer pairs verified the presence of the smaller, PE-excluded mRNA in untreated X^on^-EGFP cells, and the larger, PE-included mRNA in treated cells **(Fig. 5D-E)**. However, untreated X^on^-SALL1 cells contained an unexpected ∼700bp product, which was sequence-confirmed to be the unspliced pre-mRNA. In treated X^on^-SALL1 cells, only the expected splice product was detected **(Fig. 5E)**. When using primers flanking the PE, we detected the expected mRNAs in both untreated and treated X^on^-SALL1 cells, but failed to detect the larger unspliced pre-mRNA **(Fig. 5D)**. These results suggest that unspliced X^on^-SALL1 pre-mRNA accumulates and is translated into full-length SALL1 protein, despite the absence of the PE and start codon.

**Figure 5:**
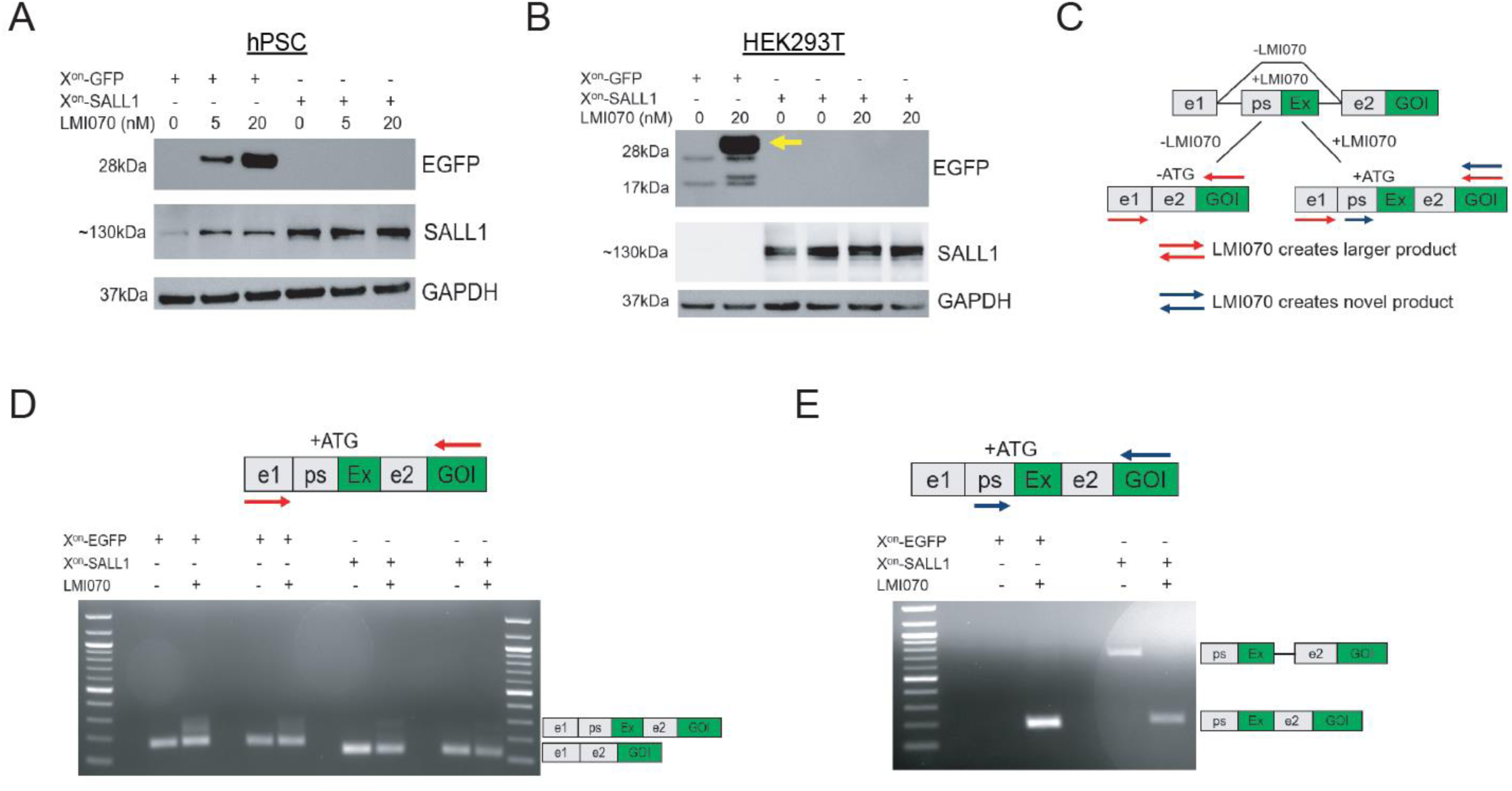
Incorporating N-terminal 2A peptides can improve inducible X^on^ systems. **(A,B)** Western blot analyses of X^on^-EGFP and X^on^-SALL1 transgenes in hPSCs (panel A) and HEK293T cells (panel B). Expected molecular weights are ∼30kDa for EGFP (yellow arrow in panel B indicates expected product), ∼140kDa for SALL1, and ∼37kDa for GAPDH. **(C)** PCR strategy for detecting X^on^ splice products. Primer pairs are represented by red and blue arrows. **(D,E)** PCR results for both primer pairs, with detected mRNAs indicated by schematics to the right of each gel. Expected product sizes for panel D are 164bp and 223bp for X^on^-EGFP with and without LMI070, and 125bp and 184bp for X^on^-SALL1. Expected product sizes for panel E are 183bp for X^on^-EGFP and 202bp for X^on^-SALL1 with LMI070. Unexpected X^on^-SALL1 product is 684bp and represents unspliced pre-mRNA.

We tested several strategies to ameliorate the apparent X^on^-SALL1 RNA processing defect, including codon optimization and addition of P2A peptides, and tested these constructs in HEK293Ts. While codon optimization modestly reduced the leaky expression, P2A peptides did not **(Fig. S6B,E)**. Surprisingly, an X^on^-EGFP-P2A-SALL1 transgene maintained EGFP inducibility despite leaky SALL1 expression, and the same effect was observed with an X^on^-EGFP-P2A-TREM2 transgene **(Fig. S6C-D)**. In contrast, both wild-type and Alzheimer’s mutant TREM2 were well controlled by LMI070 when EGFP-P2A was absent **(Fig. S6D)**. To test whether non-start ATG codons within EGFP-P2A allow for drug-independent translation, we mutated all five ATG codons (all located within EGFP) to isoleucine, but this did not prevent leakiness of either SALL1 or TREM2 **(Fig. S6B,E)**. Finally, we tested SMAD3, another candidate TF from our transposon experiments, and GRN, a secreted growth factor linked to lysosomal function and multiple neurodegenerative diseases[46]. Both genes displayed LMI070-inducible expression in HEK293Ts, and addition of an upstream P2A peptide improved both systems **(Fig. 6A-B)**. AAVS1 targeting of X^on^-P2A-GRN in hPSCs allowed for tightly controlled, tunable GRN overexpression **(Fig. 6C)**. We then differentiated these hPSCs to hMGLs and treated with LMI070, which resulted in a ∼76% increase in secreted GRN **(Fig. 6D-E)**. In summary, X^on^ allows for tunable transgene expression in hPSCs, hMacs and hMGLs, with leakiness and inducibility depending on the transgene. Addition of a 2A peptide upstream of the transgene can improve transgene control, and this system can inducibly express disease-associated genes with established therapeutic relevance in microglia-like cells[46, 47].

**Figure 6:**
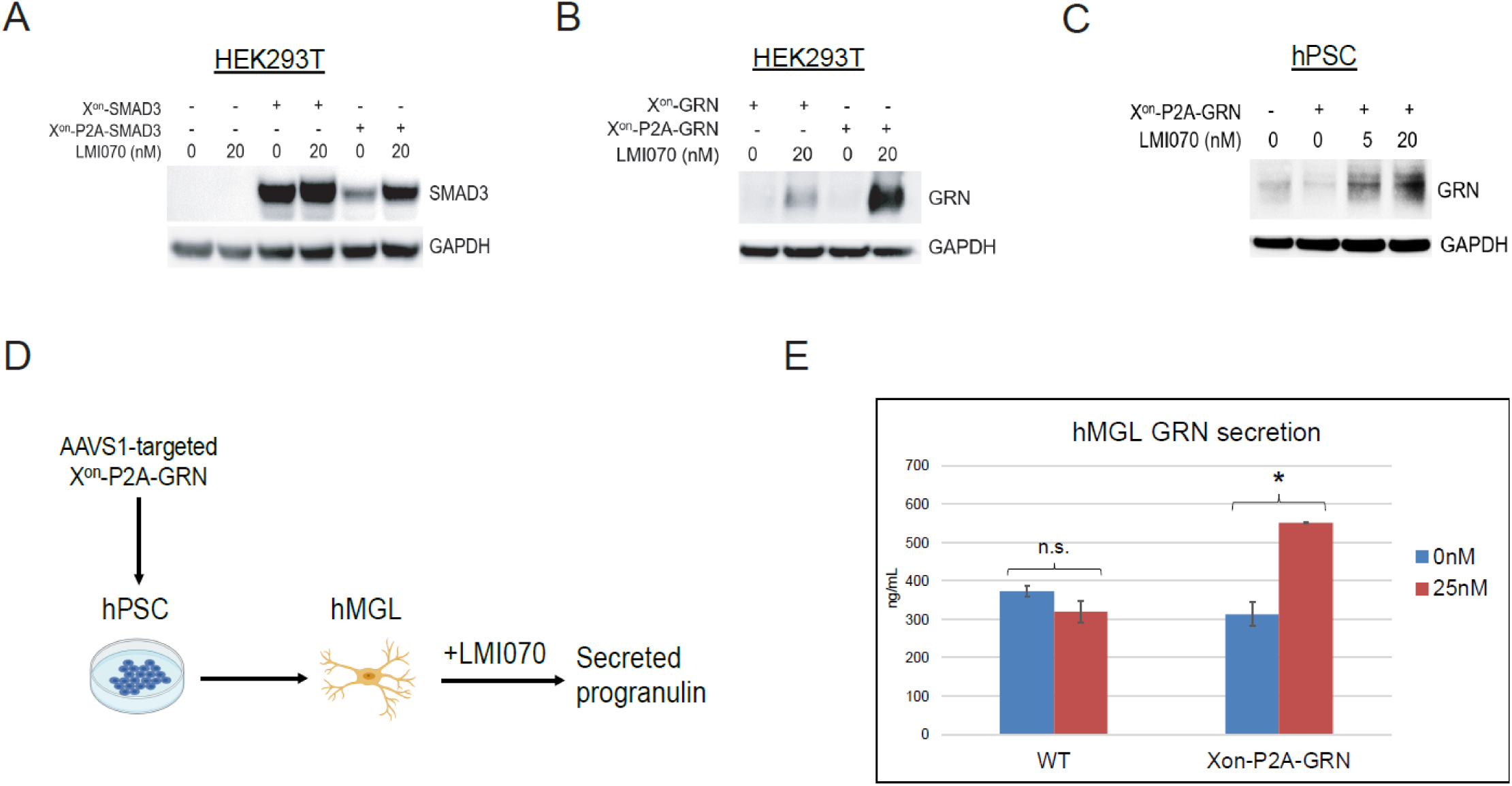
Tunable GRN overexpression with X^on^ in hPSCs and hMGLs. **(A,B)** Western blot analysis of HEK293Ts transfected with X^on^-SMAD3 and X^on^-GRN AAVS1 donor constructs. **(C)** Western blot analysis of hPSCs with AAVS1-targeted X^on^-P2A-GRN transgene. **(D)** Schematic of X^on^-mediated GRN overexpression in hMGLs. **(E)** ELISA quantification of secreted GRN from WT and X^on^-P2A-GRN hMGLs, treated with 0nM or 25nM LMI070 for 72h. n.s.=not significant; *P<0.05.

### X^on^-Cas9 genome editing performance depends on sgRNA delivery strategy

We next tested whether X^on^ can be used for inducible genome editing, and cloned Cas9 downstream of the X^on^ splicing cassette. We transduced WT and X^on^-Cas9 hPSCs with a lentivirus containing a validated sgRNA targeting the transcriptional start site (TSS) of the endogenous B2M gene[48], differentiated these cells to hMacs and hMGLs, and treated with LMI070 **(Fig. 7A)**. Using a T7 endonuclease I cleavage assay, we observed that ∼30-40% of alleles were edited in X^on^-Cas9 cells, regardless of drug treatment **(Fig. 7B)**. These results suggest that X^on^-regulated Cas9 is sufficiently leaky to achieve maximal genome editing in the absence of drug with a constitutively expressed sgRNA. To test whether transient sgRNA delivery could mitigate this, we employed a dual targeting approach with X^on^-Cas9 and an mScarlet (mSc) reporter protein inserted into each AAVS1 allele **(Fig. 7C)**. Targeted cells were transiently transfected with plasmids expressing mSc-targeting sgRNAs (pEGFP-sgRNA (3)), with or without LMI070 treatment. 48h post-transfection, we isolated transfected cells by EGFP expression, and cultured for an additional 5 days in LMI070. Only LMI070-treated cells transfected with the sgRNA plasmids showed a reduction of mSc+ cells, indicating successful induction of Cas9 editing with this approach **(Fig. 7D-E)**. These results demonstrate that while the X^on^ system can be used for Cas9-mediated genome editing in specific experimental contexts, further improvements are necessary to broaden its utility.

**Figure 7:**
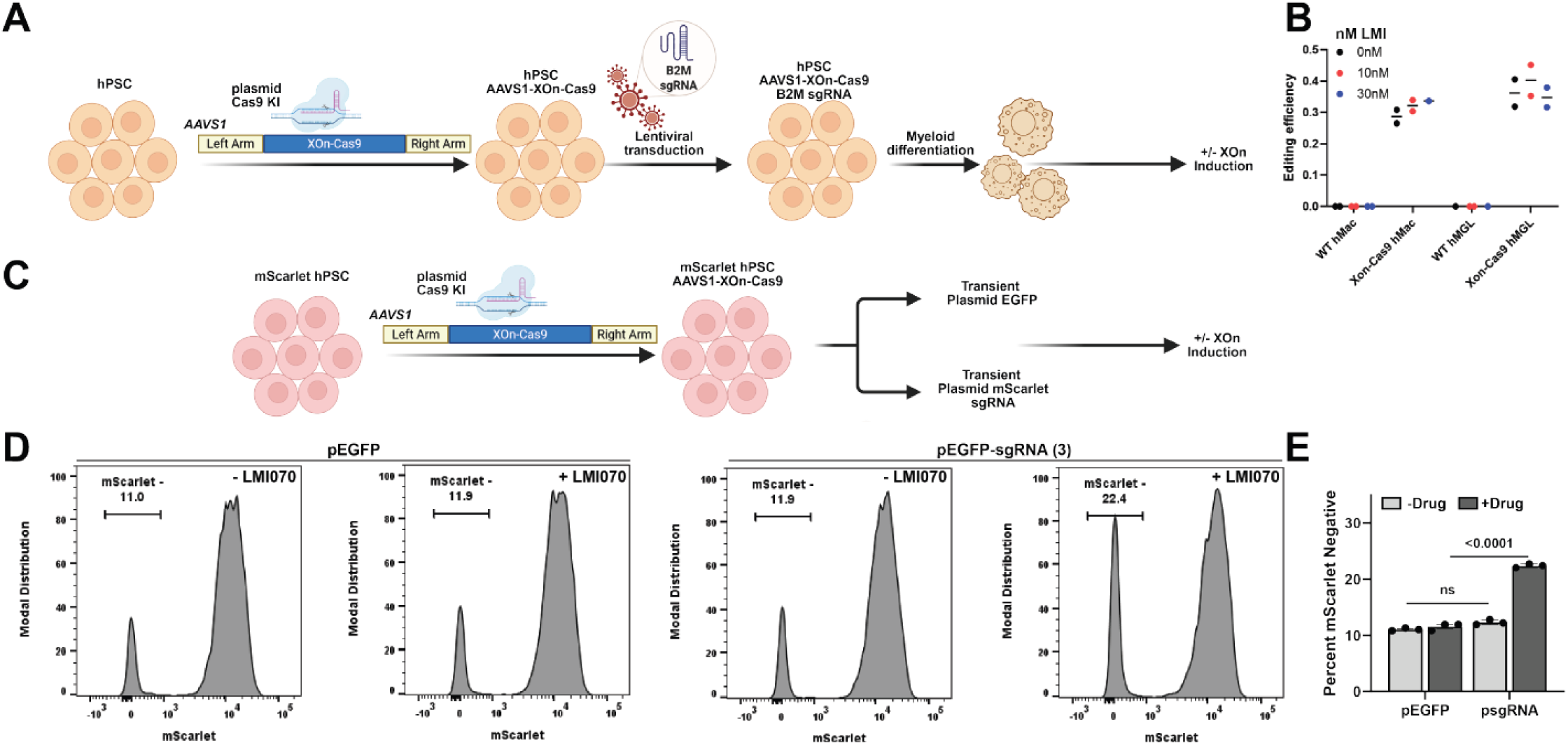
X^on^-Cas9 genome editing performance depends on sgRNA delivery strategy. **(A)** Schematic representation of high-sensitivity terminal Cas9-double strand break assay in X^on^-Cas9 hPSCs and differentiated cells with constitutive sgRNA expression. **(B)** Editing efficiency at the B2M TSS in WT and X^on^-Cas9 hMacs and hMGLs with and without LMI070 treatment. Individual circles represent biological replicates, and horizontal lines represent replicate mean values. **(C)** Schematic representation of high-sensitivity terminal Cas9-double strand break assay in X^on^-Cas9/mScarlet hPSCs with transient sgRNA expression. **(D)** Representative flow cytometry plots of X^on^-Cas9/mScarlet hPSCs with and without concurrent LMI070 induction at the time of transfection with non-sgRNA control (pEGFP) or mScarlet sgRNA (pEGFP-sgRNA (3)) plasmids. **(E)** Summary quantification of mScarlet expression in three replicate KO experiments. n.s.=not significant

## DISCUSSION

Temporal and quantitative control of transgenes are critical for determining the functions of genes on cell and organismal phenotypes. While systems providing such control are relatively advanced and well-characterized in immortalized cell lines, more physiological systems such as *in vivo* models, human primary cells and pluripotent stem cell (hPSCs)-based models still face substantial challenges in this area[6]. hPSCs are a self-renewing source of genetically modifiable cells that can be obtained fairly easily from patients, and can generate any cell type or tissue in the human body. Thus, they are ideally suited for gene function studies in human disease-relevant contexts[7]. In particular, hPSC-based systems may provide much-needed alternatives to animal models for diseases that still lack satisfactory treatments, despite decades of animal model research[18, 20, 49, 50]. Concerns have been reported about the performance of inducible transgenic systems in hPSC-derived tissues, but systematic assessments of these systems are lacking. Here we compared inducible transcriptional (Tet-On), post-translational (DHFR degron), and post-transcriptional (X^on^ alternative splicing) systems across pluripotent and differentiated myeloid cells. Using hPSC-derived macrophages (hMacs) and microglia (hMGLs) as test systems, we characterize the strengths and limitations of these systems.

The silencing of Tet-On transgenes we observe aligns with previous reports in human and mouse pluripotent stem cells[10, 11, 35, 36, 51, 52], although published results vary depending on the cell lineage, transgene, and potentially other factors such as hPSC background. As DNA methylation has been reported to underlie Tet-On silencing, we employed a similar approach as a previous study to prevent Tet-On silencing by recruiting the TET1 demethylase domain to the TRE promoter before transgene induction[35], but were not successful. More detailed characterizations of the epigenetic mechanisms underlying Tet-On silencing will be necessary for informing future modifications of these systems. While one study successfully achieved post-differentiation shRNA- and CRISPR-based gene knockdowns by inducibly expressing small RNAs with an extensively optimized Tet-On system[11], the inherent difficulties of Tet-On/Off systems may preclude broad and consistent use in hPSCs.

As we and others have demonstrated relatively stable expression of hPSC safe-harbor-targeted transgenes when driven by the CAG promoter[6, 33], systems that are inducible at post-transcriptional levels are attractive. The DHFR degron-based approach is one such system, and has been used to perform CRISPR screens in hPSC-derived neurons and microglia[39–42]. While not widely adapted as far as we know, this system is one of the only existing methods that allows for inducible gene activation or repression in fully differentiated cells. Similar to prior studies, we observe optimal transgene induction with degrons fused to both N- and C-termini, compared with single degron tags[39]. However, even the optimal degron systems perform relatively poorly in hMacs and hMGLs, calling into question the utility of this approach for expressing diverse transgenes across physiological levels. An important caveat with DHFR degrons is that fusing the ∼20kDa proteins to both termini of a protein of interest is reasonably likely to interfere with normal protein function, particularly for proteins with N-terminal signal peptides or other critical residues near the fusion site. Other degron systems such as auxin-inducible degrons, FKPB12, and molecular glues[53] warrant investigation in hPSC-based systems.

We obtained promising results with several transgenes placed under control of the X^on^ splicing switch, which is controllable by branaplam, a small molecule splicing modulator[43]. Branaplam was originally developed as a splice enhancer of exon 7 in the SMN2 gene, which increases full-length SMN2 protein and compensates for loss of SMN1 function in spinal muscular atrophy[44]. While eventually discontinued for this indication, branaplam is also a relatively specific enhancer of pseuodoexon inclusion in the human *SF3B3* gene, which led to the development of the X^on^ system in immortalized cell lines and mice[43]. The robust inducibility and tunability, requiring only low nanomolar doses, may provide an advantage over other systems with regards to off-target drug effects. For example, trimethoprim and doxycycline are typically used in the high nanomolar or micromolar ranges for DHFR-tagged and Tet-On/Off transgenes, respectively. While some studies have shown toxic or otherwise unwanted effects of branaplam human cells, these effects are not seen at low-to-mid nanomolar doses[54, 55], which were sufficient for robust transgene induction in this study. Another advantage of X^on^ is the much smaller ∼5.5kDa N-terminal tag, which can be reduced to a single proline by incorporating a 2A ribosome skipping peptide, as shown in this work. The lack of generalizability of X^on^ to transgenes such as the ∼4kb SALL1 may reflect inherent sensitivity of the branaplam-mediated splice switch to longer downstream sequences, which may interfere with splicing, transport, or other RNA processing steps[56]. Similar issues may underlie the branaplam-independent genome editing we observe in hPSCs expressing X^on^-Cas9 (also ∼4kb), with constitutively expressed sgRNA. During the writing of this manuscript, other groups have reported improvements to X^on^ by modifying sequences within or outside the pseudoexon[57, 58]. While these approaches have not, to our knowledge, been tested in fully differentiated hPSC-derived tissues, they show marked improvement over the original system in immortalized cell lines and mice. In addition, recent advances in viral transduction of myeloid cells, described by our group and several others, provide additional options for targeted transgene expression in hMacs, hMGLs, and other cell types previously not amenable to viral-based approaches (Gallagher et al, medRxiv, 2025) [59–61].

In summary, this study provides a comprehensive evaluation of inducible gene expression systems in hPSCs and differentiated myeloid cells. Our findings highlight the functional capabilities and pitfalls of these systems and establish a framework for combining and improving upon currently available systems. While no system in this study fully met the demands of no leakiness, robust induction, and tunability, the X^on^ system may represent the most tractable existing method for hPSC-based studies. Further characterization of these and other systems in other hPSC-derived lineages will complement these results, and will have important implications for the use of hPSC-based systems in basic and translational research.

## ACKNOWLEDGMENTS

We thank the flow cytometry core at the Whitehead Institute for Biomedical Research, the lab of Jonathan Weissman (Whitehead/MIT/HHMI) for the generous gift of the X^on^-EGFP piggyBac vector, and Philip A. Sharp (MIT) for helpful discussions. This work was funded by the National Institutes of Health (R01AG058002, F32AG060695, T32EB016652), Novo Nordisk, Alzheimer’s Association (AARF-22-973579) and the Wellcome Leap HOPE program.

## AUTHOR CONTRIBUTIONS

Conceptualization (M.D.G., A.S.K., D.J.M., R.J.), formal analysis (M.D.G., A.S.K., T.L., A.D., H.H.), funding acquisition (R.J., D.J.M., M.D.G., A.S.K.), investigation (M.D.G., A.S.K., Q.L., T.L., A.D., B.F.H., M.L., H.H.), methodology (M.D.G. and A.S.K.), project administration (R.J. and D.J.M.), supervision (R.J. and D.J.M.), visualization (M.D.G., A.S.K., A.D.), writing – original draft (M.D.G., A.S.K., D.J.M., R.J.), writing – review & editing (M.D.G., A.S.K., A.D., M.L., D.J.M., R.J.)

## METHODS

### Molecular cloning

Molecular cloning was carried out by inserting DNA fragments into restriction enzyme-digested backbone plasmid vectors using New England Biolabs (NEB) NEBuilder HiFi DNA Assembly Master Mix. DNA fragments were generated using a mixture of NEB Q5 High-Fidelity 2X Master Mix PCR with oligonucleotide primers synthesized from Millipore Sigma-Aldrich, Integrated DNA Technologies (IDT) synthesized double-stranded DNA fragments, and/or additional restriction enzyme-generated DNA fragments from other plasmid inserts. Detailed descriptions of insert generation are described in each construct design methods section. HiFi assembly reactions were transformed into NEB 5-alpha, 10-beta, and Stable cells, with LB agar plates incubated at 30C for 16-24h. Colonies were grown in LB with 50ug/mL ampicillin at 30C overnight, and plasmids were purified with the Omega-TEK E.Z.N.A. Plasmid DNA Mini Kit. Open reading frames (ORFs) were validated by comprehensive Sanger sequencing (Eton Bioscence) or whole plasmid Nanopore sequencing (Plasmidsaurus). Correct clones were expanded for plasmid isolation with E.Z.N.A. Endo-Free Plasmid DNA Maxi kits, which were used for transfections or cloning into lentiviral plasmids.

### Generation of transposon constructs

SALL1, TAL1 and MEF2A ORFs were PCR amplified from GenScript cDNA ORF clones. SMAD3, MEF2C and KLF2 were amplified from CS2 Flag-Smad3 (Addgene #14052), MSCV-IRES-Tomato MEF2C (Addgene #89715) and FUW-TetO-loxP-hKLF2 (Addgene #60850) plasmids. All amplicons were generated with the Roche KAPA HiFi HotStart ReadyMix PCR kit, with 5% DMSO for GC-rich templates. PCR primers contained 20bp overhangs complementary to either SfiI-digested pSBtet-GP or pSBtet-RP, P2A or T2A sequences. PCR products were verified by agarose gel electrophoresis and column purified with the Zymo Research DNA Clean & Concentrator kit. PCR reactions with unwanted products were gel purified with the Zymo Research Zymoclean Gel DNA Recovery kit before column purification. NEBuilder HiFi assembly kits were used to combine SALL1, SMAD3 and TAL1 fragments with pSBtet-GP (Addgene #60495), and MEF2A, MEF2C and KLF2 with pSBtet-RP (Addgene #60497). To replace the RPBSA promoter in pSBtet plasmids, EF1A and PGK promoters were PCR amplified from pLOX-EWgfp and PGK-H2B-mCherry plasmids, respectively. Purified products were ligated into KpnI-HF/Bsu36I-digested pSBtet-GP and pSBtet-RP with NEB T4 DNA ligase, followed by transformation, plasmid purification, and Sanger sequencing, as described above.

### Generation of Tet-On constructs

We generated four Tet-On-based systems. These constructs included the previously published 3^rd^ generation TetO3G system[62] and a combination Tet-Tight-based approach that expresses the 3^rd^ generation rtTA transactivating domain (rT fused to the VP64 transcriptional activator) and tTS (tT fused to the KRAB repressor domain) as a polycistronic cassette (pMF-TetTight, Addgene #193288)[63]. In the Tet-Tight system, the tTS was cloned downstream of the rtTA and separated by a P2A sequence. The remaining two systems were based on the Tet-Tight construct and contained N- or C-terminal fusions with the human TET1 demethylase domain. All Tet-On-based systems were synthesized as dsDNA fragments (IDT) and cloned into the AAVS1-CAG-EGFP donor construct, such that all rtTA and tTS-based transgenes were driven constitutively via the CAG promoter, downstream of the splice acceptor and puromycin selection cassette. In each of the four Tet-On systems, a seven repeat Tet-response element operator was cloned downstream of the rtTA/tTS transgene β-globin polyA sequence in reverse, with an open cloning site followed by an SV40 polyA sequence using PCR amplification of the pUCM-AAVS1-TO-hNGN2 (Addgene #105840) plasmid. HiFi assembly was performed to generate the final products, as described above. The AAVS1 CAG-driven EGFP reporter was again synthesized as a dsDNA fragment (IDT) with overhangs (left: gcgctagcacgcgtcagctgactagaggatccccgggtac, right: gtaccgagctcgaattctccaggcgatctgacggttcact) following KpnI digestion of the parent Tet-system plasmid.

### Generation of DHFR degron constructs

Synthesized dsDNA fragments (IDT) were used to generate EGFP with N-terminal, C-terminal or dual-DHFR tags, and were assembled into the AAVS1-CAG-EGFP donor construct. We used previously established trimethoprim (TMP)-responsive degron sequences[38] and a separate dual degron strategy employed in hPSC-based CRISPR screens[39–42].

### Generation of X^on^ constructs

An X^on^-GFP piggyBac plasmid was kindly gifted by Jonathan Weissman’s lab at the Whitehead Institute. We amplified the X^on^-GFP cassette from this plasmid with primers containing 20bp overhangs for HiFi assembly with AgeI-HF/EcoRV-HF-digested AAVS1-tdTomato donor construct (Addgene #159275). The resulting X^on^-GFP donor construct was used to test this transgene in hPSCs. To facilitate rapid cloning of different transgenes, an additional X^on^ donor construct (X^on^ “open”) was generated with an AflII cut site for cloning in a gene of interest, followed by a subsequent round of HiFi assembly with synthesized DNA fragments containing plasmid homology arms: AATTGTTGTTTCCCGTGGGTAAGAGGATCC (left) and AATTGTTGTTTCCCGTGGGTAAGAGGATCC (right). For specific transgenes, (EGFP, SALL1, SMAD3, TREM2, GRN, Cas9) the corresponding fragments were synthesized without the start codon, but with 5’ and 3’ overhangs to allow for assembly into the digested donor construct. The R47H TREM2 mutation was tested due to its association with Alzheimer’s disease risk[64]. P2A sequences were added by incorporating the necessary sequences into the overhangs, and EGFP-P2A was added upstream of the genes of interest with a separate synthesized dsDNA fragment. All transgenes were cloned into either the X^on^ “open” donor or the original AAVS1-tdTomato-based donor.

### Generation of sgRNA constructs

For transient mScarlet sgRNA experiments, the mScarlet sgRNA was cloned into a modified px330-Cas9 plasmid in which Cas9 was replaced with EGFP (px330-EGFP), such that only sgRNA and EGFP would be expressed in transfected cells. sgRNAs were inserted into px330-EGFP using the BbsI-HF restriction enzyme, as previously described[65].

### HEK293T cell culture and transfections

HEK293Ts were cultured in DMEM supplemented with 10% inactivated cosmic calf serum, 2mM L-glutamine and 1% penicillin/streptomycin. Cells were passaged every 2-3 days with trypsin at a 1:8 split ratio. Transfections were performed with X-tremeGENE 9 reagent, with up to 4ug of total DNA per well of a 6 well plate, and 3uL of X-tremeGENE 9 per 1ug of DNA.

### hPSC cell culture and transfections

Previously published female iPSCs[22] and H1 ESCs were cultured and passaged as previously described (Gallagher et al, medRxiv, 2025). Transfections were performed with ViaFect (Promega), with up to 4ug of total DNA per well of a 6 well plate, and 2uL of ViaFect per 1ug of DNA. ViaFect and DNA were diluted in Opti-MEM, combined, and added to cells with E8 media in a total volume of 2-3mL per well of a 6 well plate. For cells maintained in mTeSR Plus or StemFlex, E8 was replaced with maintenance media the next morning. For promoter testing, pmax-GFP, PGK-H2B-GFP, pLOX-Ewgfp (Addgene #12242), pcag-GFP (Addgene #11150) were transfected into H1 ESCs and assessed by FACS or fluorescence microscopy.

### HEK293T transposon experiments

Cells were transfected with pSBtet-based transposons and pCMV(CAT)T7-SB100 (Addgene #34879), which encodes the SB100X transposase. Mass ratios of 1:3 to 1:20 were tested, with 1:20 producing the largest number of integrations. GFP+ or tdT+ cells were isolated by FACS 7-10 days after transfection, and these populations were frozen in liquid nitrogen and treated with 0.75-7.5ug/mL doxycycline (dox) to assess dox-inducible expression of transcription factors. Cell lysates were extracted 2-4 days after dox treatment for Western blot analysis. For generation of cells with integrations of both pSBtet-GP-TF and pSBtet-RP-TF transposons, cells were co-transfected with both transposon constructs and SB100X.

### hPSC transposon experiments

iPSCs were transfected with pSBtet-based constructs and pCMV(CAT)T7-SB100 using the ViaFect transfection protocol described above. As with HEK293Ts, excess SB100X was used to maximize integration efficiency, and FACS-based isolation of cells with integrations was performed 7-10 days after transfection. Dox was added at 0.2 or 2ug/mL to iPSCs or iPSC-derived myeloid progenitors for 2-4 days before lysate extraction for Western blotting. Puro resistance was assessed by adding 2ug/mL puro to pSBtet-GP-TF myeloid progenitors pSBtet-EF1A-GFP iPSCs before and throughout hMGL differentiation.

### hMac and hMGL differentiation

hMacs and hMGLs were generated with two differentiation protocols, one using an embryoid body approach, and one using a monolayer approach, as previously described[22, 66], (Gallagher et al, medRxiv, 2025).

### hPSC AAVS1 targeting

To generate the stable expression of constitutive or inducible transgene reporters, our previously published AAVS1-CAG-EGFP donor construct[33] was used as a starting template. Genome targeting was performed as previously described using PX458-AAVS1-sg (Addgene #194721) to express Cas9 and an AAVS1*-*targeting sgRNA (GGGGCCACTAGGGACAGGAT)[34], and the various AAVS1 donor constructs generated in this manuscript. Introduction into cells was either carried out via electroporation (Lonza 4D-Nucleofector System, according to the manufacturer’s instruction) or transfection. Transfections were performed with ViaFect, using 2.67ug of donor construct and 1.33ug of PX458-AAVS1-sg per well of a 6 well plate. For transgenes with puromycin (puro) resistance, cells were treated with sequentially increasing doses of puro (typically 0.5, 1 and 2ug/mL) for 2 days each, and surviving colonies were frozen in liquid nitrogen and maintained without subcloning for subsequent experiments. 10mM Y-27632 was typically needed for the first two days of puro treatment to prevent transgenic cells from listing off in the presence of widespread death.

### Tet-On experiments

To compare the different Tet-On design strategies, we targeted H1 hPSCs at the AAVS1 locus with the various Tet-On systems described above. After puro-based selection, we performed a time course of induction in hPSCs using 1µg/mL of doxycycline in mTeSR Plus medium, and assessed EGFP fluorescence by flow cytometry. Measurements of induction efficiency were taken at 48, 96, and 144h post-treatment, with fresh dox-supplemented media replaced at each timepoint. For silencing experiments, hPSCs were differentiated into hMacs without prior dox exposure. hMacs were then subjected to 1µg/mL doxycycline for 144h and EGFP fluorescence was assessed by flow cytometry.

### DHFR degron experiments

To compare performance of the different degron designs, we targeted H1 hPSCs at the AAVS1 locus with the above degron constructs as described above. We then performed a trimethoprim (TMP) (Thermo Fisher Scientific, cat. #J67175.XF) dose-response curve via serial dilution of 2.5µg/ml (8.6 µM) to 0.312 µg/mL (1.1µM) diluted in mTeSR Plus medium. 24h post-treatment, EGFP intensity was measured via flow cytometry and compared that of WT (non-engineered) hPSCs, as well as each engineered degron without TMP treatment. For silencing outcome comparisons, the engineered H1 hPSCs were differentiated into hMacs and treated with 2.5µM TMP for 24h, with similar flow cytometry-based assessment of induction capacity.

### X^on^ transgene induction experiments

To evaluate X^on^-EGFP in hPSCs, we performed a LMI070 (MedChemExpress, cat #HY-19620) dose-response curve via dilution of 100nM to 2nM in mTeSR Plus medium. After 24-48h, we assessed fluorescence intensity via flow cytometry and/or Western blot, and compared to WT (non-engineered) hPSCs untreated (DMSO only) X^on^-EGFP cells. For differentiated cells, dose response curves were performed after differentiation of X^on^-EGFP hPSCs to hMGLs and hMacs, using both embryoid body[66] and monolayer[22] protocols. GFP was assessed by flow cytometry after 24h of treatment. X^on^-EGFP, X^on^-SALL1, X^on^-TREM2, X^on^-SMAD3 and X^on^-GRN, including variations of these transgenes (see Fig. S3, 5 and 6), were tested by transfecting HEK293Ts with AAVS1 donor constructs containing each indicated transgene, followed by treatment with 20nM LMI070 or DMSO for 24h-48h. Western blots were used to assess expression of the transgenes. X^on^-SALL1 and X^on^-P2A-GRN were additionally targeted to AASVS1 in hPSCs, and tested for inducibility in hPSC-derived myeloid progenitors and hMGLs made with the monolayer hMGL protocol. X^on^-SALL1 hPSCs and myeloid progenitors were treated with 5-20nM LMI070 or DMSO for 24h-48h, and SALL1 expression was determined by Western blot. X^on^-P2A-GRN hMGLs were treated with 25nM LMI070 or DMSO for 72h, and levels of secreted GRN were determined by ELISA. In all experiments, WT cells lacking X^on^ transgenes were treated with the same LMI070 or DMSO doses as controls. For the GRN ELISA experiments, X^on^-P2A-GFP hPSCs were differentiated into hMGLs with the monolayer protocol. Myeloid progenitors were plated at 1 million cells in 4mL microglia maturation media per well of a 6 well plate, with two biological replicates per condition. Five days later cells were treated with DMSO or 25nM LMI070, and the media was collected three days afterwards. Media from each well was split into two 1.5mL tubes, filtered through a 0.45um mesh filter, and frozen in 1 mL aliquots at −80C.

### Integrated sgRNA X^on^ Cas9 induction experiments

To generate the AAVS1 X^on^-Cas9 targeting construct, a spCas9 dsDNA fragment, minus a start codon, was synthesized and cloned into the AAVS1 X^on^-Open construct, and targeted to AAVS1 in hPSCs, as described above. The AAVS1 X^on^-Cas9 donor template contains a promoter-less neomycin resistance cassette upstream of the CAG promoter, with a splice acceptor for endogenous expression of the selection marker. sgRNA lentivirus was made as previously described (Gallagher et al, medRxiv, 2025). Briefly, HEK293Ts were transfected with pBA904 BFP-expressing Perturb-seq plasmid (Addgene #122238) containing a validated B2M-targeting sgRNA, pCMV-VSV-G and psPAX2-chp6, followed by virus collection 48h-72h later. WT and X^on^-Cas9 hPSCs cultured in 300uL mTeSR Plus in a 24 well matrigel-coated plate were transduced with 120uL of B2M lentivirus, and 300uL IMDM was added to bring the total volume in each well to 600uL. Cells received a full mTeSR media change the next day, and were passaged and expanded one day later. Transduced cells were selected with puro, frozen in liquid nitrogen, and differentiated to hMacs and hMGLs with the embryoid body protocol. After 7 days in hMac or hMGL maturation media, cells were treated with 0, 10 or 30nM LMI070, then treated a second time with fresh media 3 days later. Two days after the second treatment, BFP expression was analyzed by flow cytometry and gDNA was purified with the QIAgen DNeasy Blood & Tissue kit. gDNA concentration was determined by QuBit, and Cas9 editing efficiency was quantified by T7 endonuclease I digestion of heteroduplexes with the NEB EnGen Mutation Detection Kit. Digested PCR products were run on a bioanalyzer to determine the ratio of edited to total product. Data were analyzed and plotted with GraphPad Prism.

### Transient sgRNA X^on^ Cas9 induction experiments

X^on^-Cas9 hPSCs were targeted with a constitutively active CAG-mScarlet-3XNLS transgene into the AAVS1 locus, containing with a splice-acceptor driven puromycin-resistant cassette for selection. Dual puro/neo antibiotic selection yielded a population of inducible Cas9/mScarlet fluorescent hPSCs. For transient sgRNA expression, these hPSCs were transfected with either the modified px330-EGFP plasmid (lacking Cas9 and sgRNA) or three similar plasmids containing different sgRNAs targeting near the mScarlet chromophore residue (sgRNA #1: ATATCCCAGCTAAACGGCAG, sgRNA #2: CGGCTGCCATACATAAACTG, sgRNA #4: CTGAGCCCGCAGTTTATGTA). Cells were transfected concurrently with or without 20nM LMI070, and we performed FACS to isolate EGFP+ cells after 48h to isolate sgRNA-expressing cells. 5 days post-FACS and 7 days post-transfection, these cells were subjected to flow cytometry to assess mScarlet expression.

### Western blotting

Cell lysates were extracted with RIPA, and protein concentrations were quantified with the Pierce BCA Protein Assay kit. Lysates were denatured at 95C for 10 min, loaded onto 4-12% and 10% NuPAGE Bis-Tris Mini protein gels with 1X NuPAGE LDS Sample Buffer, and run at 100V in XCell SureLock Mini-Cell electrophoresis units. Protein was transferred from the gels to PVDF membranes in Bio-Rad Mini Trans-Blot Cells at 350mA for 2.5h. Blots were rinsed in TBST and TBS, blocked with 5% non-fat dry milk in TBST, and incubated with primary antibodies diluted in 5% milk with 0.02% sodium azide. Primary antibody incubations were performed overnight at 4C on a shaker. Secondary antibodies (Sigma-Aldrich goat anti-rabbit, goat anti-mouse and rabbit anti-goat IgG H&L Chain Specific Peroxidase Conjugate) were applied to blots at 1:10,000 dilutions in 5% milk for 1.5-2h at room temperature. Blots were developed with Western ECL or Western PicoPlus reagents either on X-ray films or with a Bio-Rad ChemiDoc Imaging System. SeeBlue Plus2 Pre-Stained Protein Standard and Precision Plus Protein Dual Color Standard were used as markers. Primary antibodies and dilutions included: abcam ab125247 anti-GAPDH (1:10,000), abcam ab6556 anti-GFP (1:5000), R&D AF2420 anti-GRN (1.25ug/mL), Novus Biologicals NBP2-61812 anti-KLF2 (1:365), Santa Cruz Biotechnology sc-17785 anti-MEF2A (1:1000), Sigma HPA005533 anti-MEF2C (1:250-1:1000), R&D PP-K9814-00 anti-SALL1 (1:1000), abcam ab40854 anti-SMAD3 (1:5000), Thermo Fisher Scientific PA5-46840 anti-TAL1 (1:1000), Cell Signaling Technology #91068S anti-TREM2 (1:1000).

### Flow cytometry

HEK293Ts were dissociated with Trypsin, hPSCs with ReLeSR or Versene, hMGLs with Trypsin (if attached) or by collection and pelleting (if floating), and hMacs collected with Versene and cell scraping. Cells were washed in FACS buffer (PBS with 1% HyClone FBS, 2mM EDTA and 25mM HEPES), stained with 0.2ug/mL DAPI and analyzed and sorted with a BD FACSAria II or BD LSRFortessa. Tet-On induction time course experiments were calibrated from day to day using Rainbow Calibration Beads (Thermo Fisher). Sorted hPSCs were plated in maintenance media containing Stem Cell Technologies CloneR or CloneR2 to facilitate survival of single cells. All flow cytometry data were analyzed with FlowJo and GraphPad Prism.

### Microscopy

All images were taken with a Nikon Eclipse Ti-U or Nikon Eclipse TE2000-U microscope and analyzed with ImageJ.

### PCR analysis of X^on^ splicing events

PCR primers were designed to detect amplicons that increase in size (pair 1), or are unique (pair 2), when the pseudoexon is spliced into the mRNA. X^on^-GFP primer pair 1 (F: GCTTTGCCATTCTTGGAAAC, R: cagacacgctgaacttgt), X^on^-GFP primer pair 2 (F: GAGTCAATCCAAGTgccAc, R: cagacacgctgaacttgt), X^on^-SALL1 primer pair 1 (F: CCACTGGCATCAGCTTTG, R: cggattggaaatgttgaggc), X^on^-SALL1 primer pair 2 (F: GAGTCAATCCAAGTgccAc, R: ttagtagggcgactcggt). RNA was extracted from X^on^-GFP and X^on^-SALL1 hPSCs 48h after treatment with 20nM LMI070 or DMSO, using the QIAGEN miRNeasy Mini Kit. 1ug of each RNA sample was DNase-treated (Thermo Fisher Scientific, cat. #EN0521), and cDNAs were generated with Quantabio qScript cDNA SuperMix (cat. #95048-025), according to manufacturer instructions. Primers were tested for efficacy and specificity by qPCR on an Applied Biosystems QuantStudio 6 Flex, using Fast SYBR Green 2X master mix (Thermo Fisher Scientific, cat. #4385616). Amplicons were then generated with NEB Q5 PCR master mix and analyzed by agarose gel electrophoresis, with NEB 100bp DNA ladder. Identify of amplicons was confirmed by Sanger sequencing (Eton Bioscience) after column purification.

### GRN ELISA

GRN ELISA was performed with the AdipoGen Life Sciences Progranulin (human) ELISA kit (cat. # AG-45A-0018YEK-KI01), according to the manufacturer’s instructions. Frozen media from LMI070-treated X^on^-P2A-GRN hMGLs was thawed and diluted 1:50 in microglia maturation media before performing the ELISA. Each frozen media aliquot from the same well were analyzed, but were averaged and not considered as biological replicates. Biological replicates were different wells of the same cell population treated with the same LMI070 dose.

### Statistical analyses

Simple linear regressions were performed to test for correlations between LMI070 dose and EGFP expression in X^on^-EGFP hPSCs, hMacs and hMGLs. R^2^ and p-values are reported in the results section. Effects of Tet-On and DHFR degron transgenes on leaky EGFP expression were analyzed with 1-way ANOVAs. Effects of the X^on^-EGFP transgene on leaky EGFP expression were analyzed with 2-tailed t-tests. GRN ELISA results were analyzed with two-tailed t-tests to determine effects of LMI070 dose on GRN levels in hMGLs. X^on^-Cas9 editing efficiency in treated and untreated B2M sgRNA-expressing hMGLs and hMacs was analyzed with a 2-way ANOVA. mScarlet knockout in X^on^-Cas9 hPSCs transfected with mScarlet-targeting sgRNAs was analyzed with a 1-way ANOVA. All statistical analyses were performed in GraphPad Prism.

## SUPPLEMENTARY FIGURE LEGENDS

**Figure S1:**
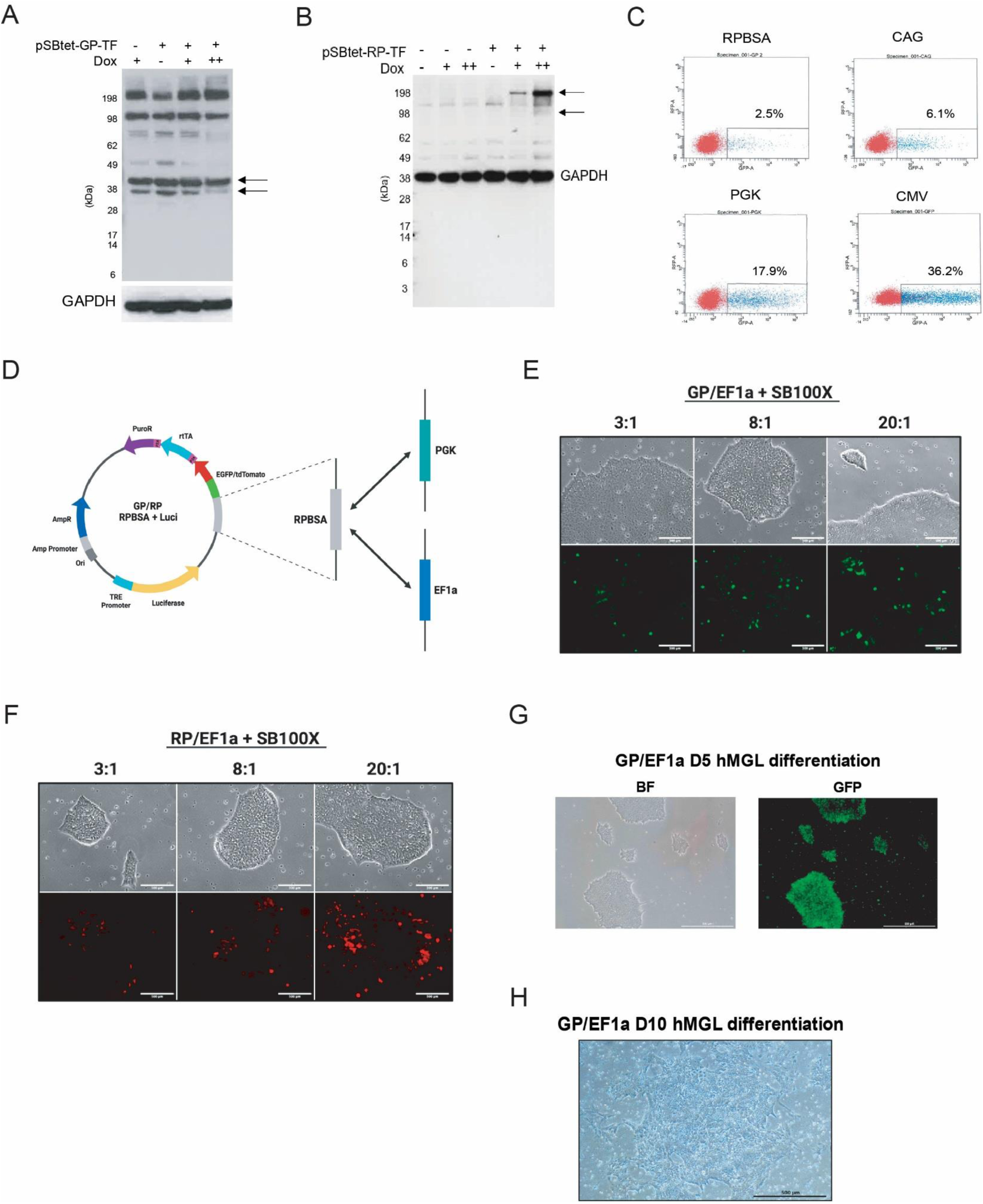
Effects of promoter replacement on pSBtet transposons in hPSCs and hPSC differentiation. **(A,B)** Western blot analyses of HEK293Ts transfected with pSBtet-GP-TF (panel A) and pSBtet-RP-TF (panel B). Panel A shows blot results for TAL1, with arrows indicating bands corresponding to the expected molecular weight of TAL1 (35-40kDa). Panel B shows blot results for KLF2, with arrow indicating dox-specific bands. As with Figure 1, dox doses are 0.75 and 7.5ug/mL. **(C)** Flow cytometry analysis of hPSCs transfected with plasmids expressing EGFP under control of RPBSA, CAG, PGK and CMV promoters. GFP+ cells are shown in blue, with the proportions indicated above the GFP+ gates. **(D)** Schematic of promoter pSBtet promoter replacement strategy. **(E,F)** Brightfield (top) and GFP (bottom) channels of hPSCs with stable integrations of pSBtet-GP-EF1A (panel E) and pSBtet-RP-EF1A (panel F). Ratios above images indicate transfection mass ratios of SB100X transposase and pSBtet transposons, with higher ratios improving integration efficiency. Scale bars are 500µm. **(G,H)** Brightfield and GFP channels from images of pSBtet-GP-EF1A hPSCs at days 5 (panel G) and 10 (panel H) of hMGL differentiation. Puro-induced cell death is visible at day 10 but not day 5. Scale bars are 500µm.

**Figure S2:**
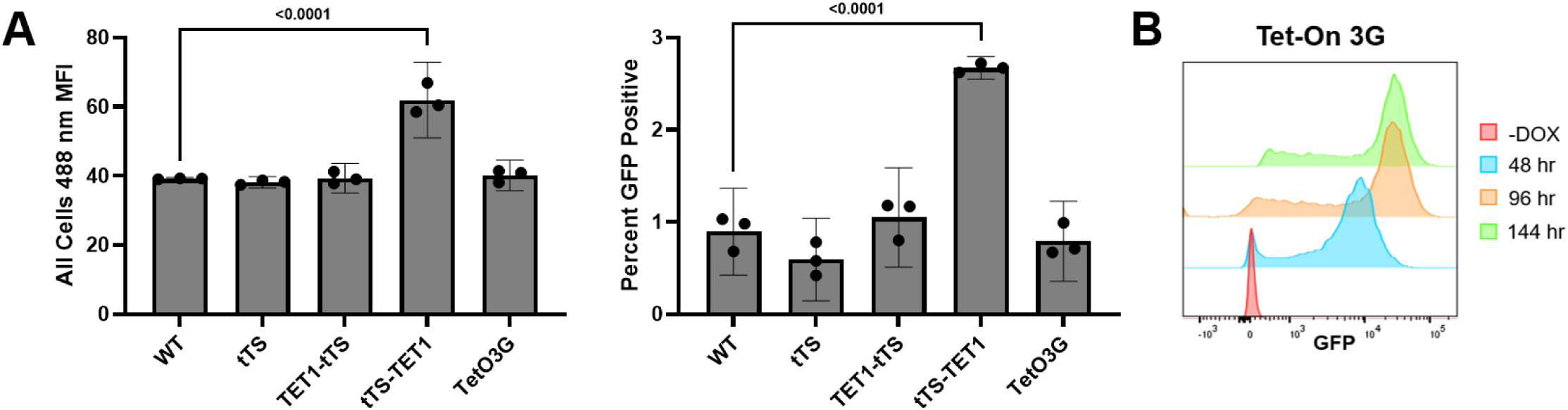
Quantification of leaky expression in Tet-On and tTS-based systems. **(A)** Geometric mean fluorescence intensity (MFI) at 488nm of all singlet gated hPSCs (left) and the percentage of EGFP+ cells when gated at ∼0.9% against unengineered WT cells, all in the absence of doxycycline (DOX). **(B)** Time course of EGFP expression response represented in hours post-DOX exposure in hPSCs engineered with the standard third-generation Tet-On system, TetO3G.

**Figure S3:**
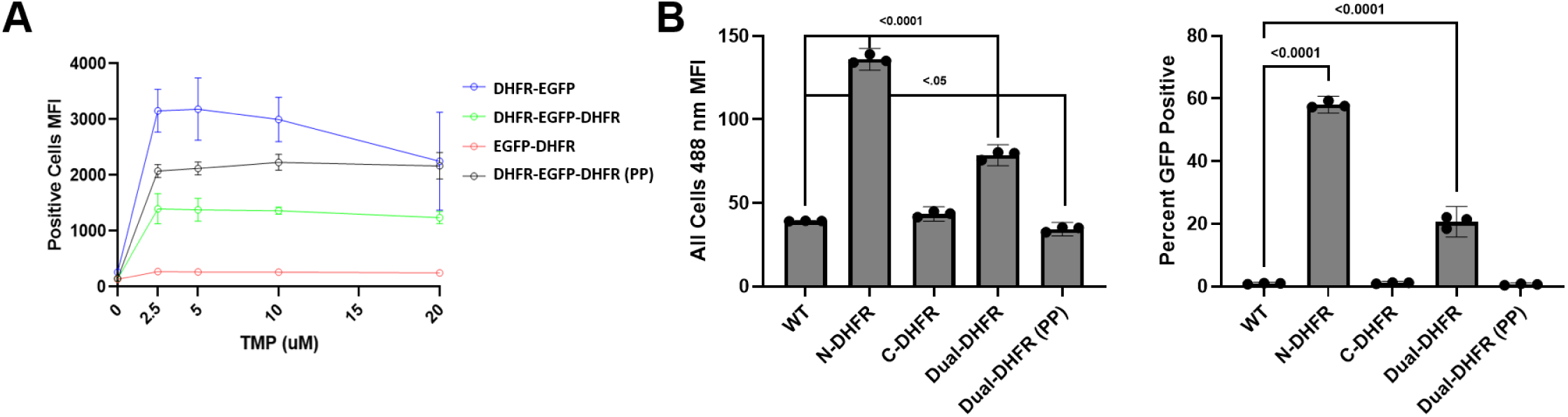
Quantification of leaky expression in DHFR degron systems. **(A)** Dose-response of EGFP expression represented 24h post-trimethoprim (TMP) exposure in hPSCs engineered with all DHFR degron systems. **(B)** Geometric mean fluorescence intensity (MFI) at 488nm of all singlet gated hPSCs (left) and the percentage of EGFP+ cells when gated at ∼0.9% against unengineered WT cells, all in the absence of TMP.

**Figure S4:**
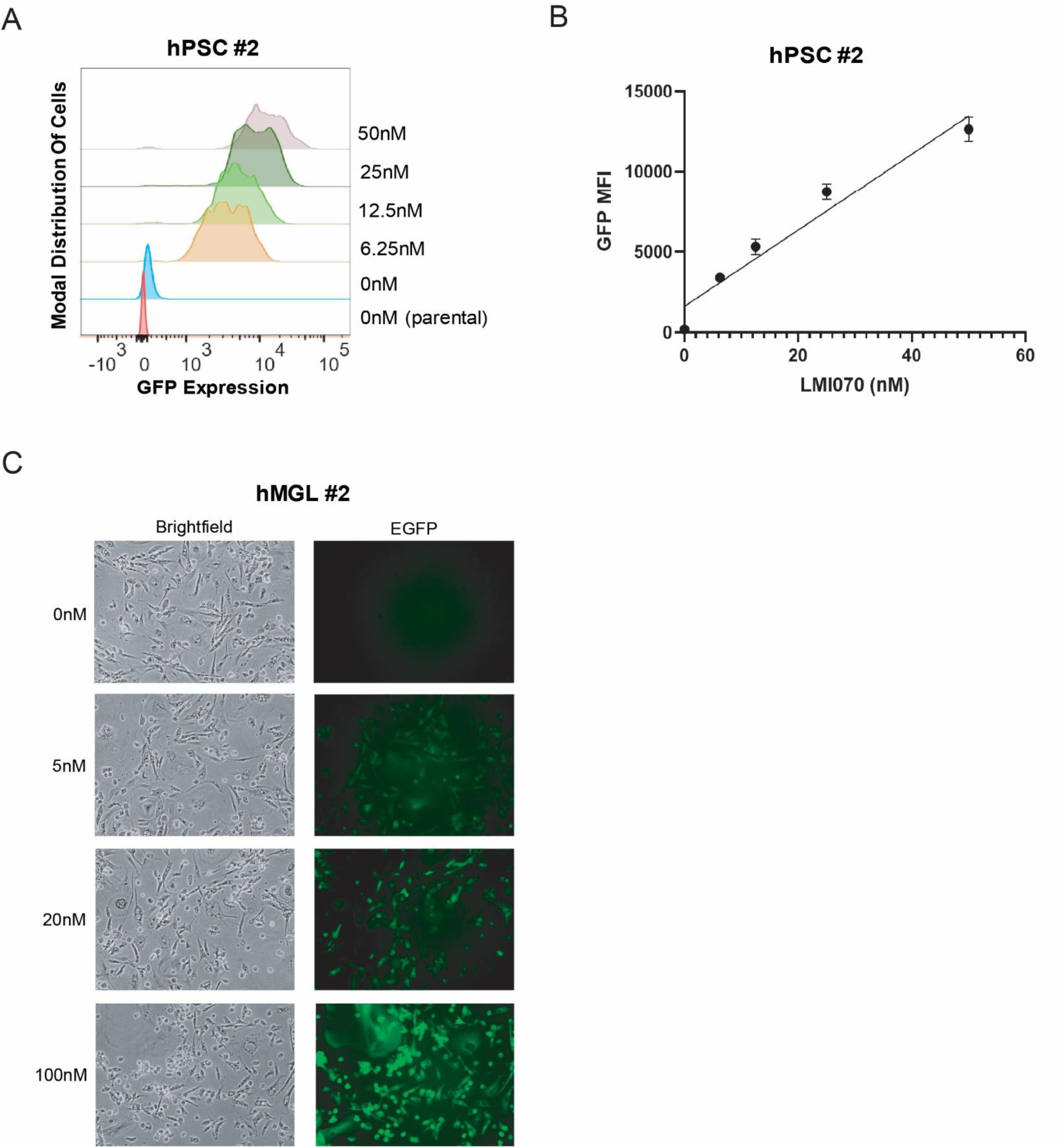
X^on^-EGFP performance in alternative hPSCs and hMGLs generated with a different differentiation protocol. **(A-B)** hPSCs from a different genetic background as those in Fig. 4 display tunable LMI070-mediated EGFP expression. **(C)** hMGLs produced from hPSCs shown in panel A, but with a different differentiation protocol[66] than those shown in Fig. 4, display LMI070 dose-dependent EGFP expression, with no obvious toxicity.

**Figure S5:**
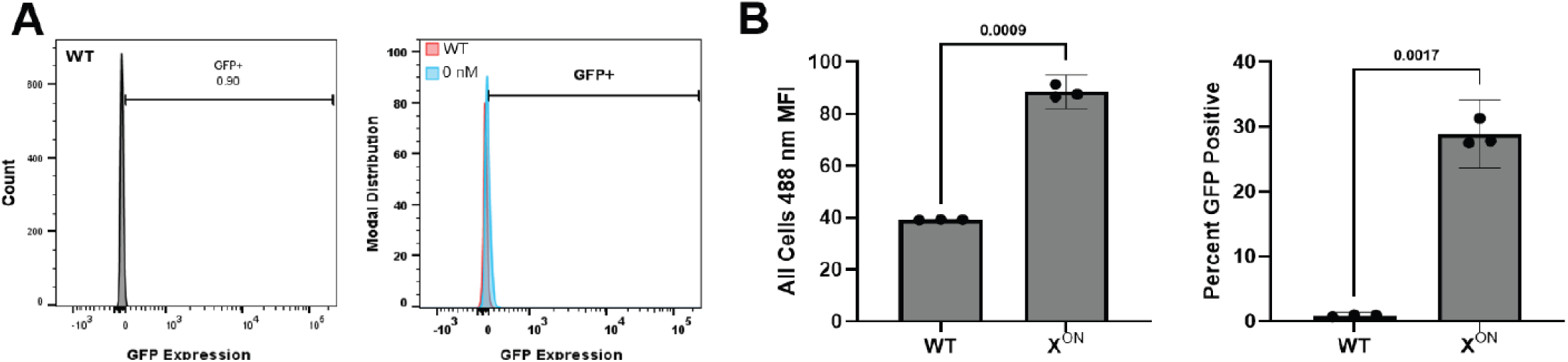
Quantification of leaky expression in X^on^ system. **(A)** Representative flow cytometry plot demonstrating distribution of 488nm fluorescence in unengineered WT hPSCs (left), overlayed with X^on^-EGFP-engineered hPSCs (right), both with ∼0.9% stringency gate illustrated. **(B)** Geometric mean fluorescence intensity (MFI) at 488nm of all singlet gated X^on^-EGFP hPSCs (left) and the percentage of EGFP+ cells when gated at ∼0.9% against unengineered WT cells, all in the absence of LMI070.

**Figure S6:**
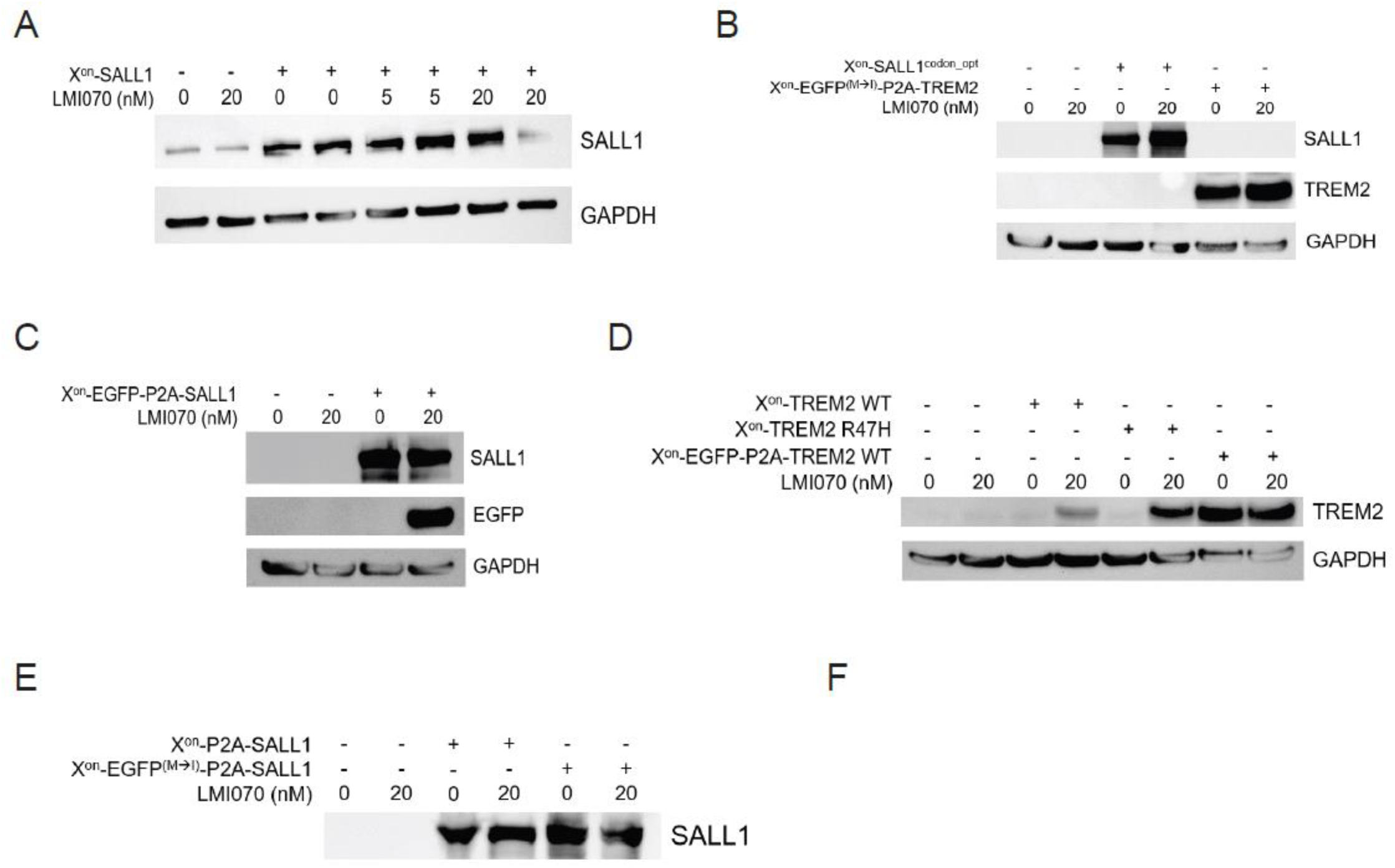
Approaches to ameliorate technical limitations of the X^on^ system. **(A)** Western blot analysis of AAVS1-targeted X^on^-SALL1 in hPSCs from a different genetic background than those shown in Fig. 5A, indicating leakiness and poor inducibility in the presence of LMI070. **(B)** Western blot analysis of HEK293Ts transfected with codon-optimized X^on^-SALL1 and Met→Ile mutated X^on^-EGFP-P2A-TREM donor constructs. **(C)** Western blot analysis of HEK293Ts transfected with X^on^-EGFP-P2A-SALL1 donor construct. **(D)** Western blot analysis of HEK293Ts transfected with X^on^-TREM2 WT, X^on^-TREM2 R47H, and X^on^-EGFP-P2A-TREM2 donor constructs. **(E)** Western blot analysis of HEK293Ts transfected with X^on^-P2A-SALL1 and Met→Ile mutated X^on^-EGFP-P2A-SALL1 donor constructs.

